# Self-generated oxygen limitation drives coral outcomes during marine heatwaves

**DOI:** 10.64898/2026.10.08.757541

**Authors:** Yusuf C. El-Khaled, Hamid H. Khan, Shannon G. Klein, Neus Garcias-Bonet, Dilip Maity, Sandip Dighe, Timofey Mukha, Abhijit K. Kushwaha, Fauzia Wardani, Mourad Oulghelou, Max S. Dhillon, Spencer Truman, Francisca C. García, Maggie D. Johnson, Michael Fox, Carlos M. Duarte, Susana Carvalho, David J. Suggett, Raquel S. Peixoto, Matteo Parsani, Tadd T. Truscott

## Abstract

Marine heatwaves cause massive, but often patchy coral bleaching. Why neighboring colonies often respond differently remains unresolved. Here, we show that interactions between metabolic demand and flow-mediated resupply generate heterogeneous oxygen microenvironments inside branching corals under heat stress. During the 2023 and 2024 Red Sea heatwaves, high-frequency *in situ* measurements revealed hypoxic to near-anoxic interbranch conditions despite well-oxygenated surrounding seawater. In 2023, neighboring colonies developed divergent oxygen regimes preceding paling and bleaching; in 2024, stronger and earlier thermal stress was associated with recurrent nighttime near-anoxia before tissue disintegration and mortality. A physics-based three-dimensional fluid-metabolism model reproduced these patterns. Oxygen depletion and coral impacts emerged when metabolic oxygen demand exceeded flow-mediated resupply. Organism-scale variability in oxygen regimes may explain divergent coral fate under shared heatwave exposure.

## Main

Marine heatwaves and extreme climate events are driving mass mortality across organisms and ecosystems worldwide ^1^, including seagrass meadows ^2^, fish populations ^3^, mangrove forests ^*4*^ and coral reefs ^5–7^. Sessile organisms are particularly vulnerable to such events due to their inability to escape deteriorating conditions, yet their biological responses are often strikingly heterogeneous despite apparent homogeneous environmental exposure ^2, 8, 9^. This disparity reveals a central limitation of current marine climate-risk frameworks: broad-scale environmental metrics identify where extreme conditions occur, but cannot explain why adjacent organisms, or even different parts of the same individual, cross physiological thresholds at different times or avoid them altogether. In scleractinian corals, this unresolved heterogeneity is directly visible as spatially patchy paling, bleaching, and tissue loss during marine heatwaves ^10^.

Patchiness in coral bleaching has been attributed to intrinsic biological variation, including differences in host traits, symbiont identity, and acclimatization history ^11, 12^. However, the inherent spatial heterogeneity of such patchiness indicates that intrinsic biological variation alone is insufficient to explain divergent bleaching outcomes under similar far-field thermal exposure. Rather, this heterogeneity points to micro-scale processes that can vary significantly at small spatial scales, such as fine-scale temperature variability ^13^ or local oxygen availability ^14^. Yet, operational bleaching-risk frameworks remain centred on broad-scale temperature metrics and thresholds ^15^ and do not incorporate oxygen dynamics, despite knowledge that oxygen availability influences coral stress from organismal ^16^ to ecosystem scales ^17^. In particular, deoxygenation can lower coral bleaching thresholds ^18^, and acute hypoxia driven by marine heatwaves can cause reef-scale bleaching, tissue loss, and mortality ^17^. Hence, it becomes apparent that the local balance between metabolic oxygen demand and flow-mediated oxygen resupply within colony microenvironments needs to be incorporated into operational bleaching-risk metrics.

Heterogeneous bleaching responses may arise when local differences in flow-mediated oxygen resupply determine whether temperature-enhanced metabolic demand produces oxygen limitation within colony microenvironments. Warming promotes oxygen depletion ^*19, 20*^ by reducing oxygen solubility ^*21*^, increasing respiratory demand ^*22*^, inhibiting photosynthetic oxygen production ^*23*^ and impairing ciliary effects that contribute to microscale ventilation ^*24, 25*^. Where fluid transport supply and ciliary ventilation cannot compensate for oxygen depletion under rising metabolic demand, oxygen conditions within organismal microenvironments may diverge sharply from those in the surrounding water. Laboratory experiments have demonstrated that corals under thermal stress can develop such localized oxygen depletion despite being submerged in oxygenated bulk water ^24, 25^. Whether such constraints develop *in situ* during marine heatwaves and contribute to coral bleaching and mortality remains unresolved.

Branching corals provide a tractable system to resolve this organism-scale gap because their three-dimensional morphology interacts with flow to create spatially heterogeneous interbranch microenvironments. Within these spaces, advective exchange, turbulent mixing, and molecular diffusion driven by gradients generated by coral metabolism regulate the transport of dissolved gases between the colony interior and surrounding reef water ^26–30^. Reduced transport can therefore allow metabolic oxygen consumption to exceed replenishment, particularly at night or during bleaching, when photosynthetic oxygen production ceases or is impaired, respectively. This mechanism may help explain why elevated water flow has been associated with reduced bleaching susceptibility and faster recovery ^31–33^.

Here, we propose a transport-metabolism framework in which bleaching susceptibility emerges from imbalances between temperature-driven metabolic demand and flow-mediated oxygen supply within coral colony microenvironments (Fig. 1). Under moderate thermal stress, local hydrodynamic differences may determine whether colonies maintain stable oxygen budgets or develop progressive oxygen depletion. Under stronger thermal stress, however, metabolic demand may exceed the buffering capacity of local transport, causing oxygen instability to become widespread. This framework predicts that increasing heatwave intensity shifts colony-scale microenvironments from stable normoxia towards increasingly unstable oxygen regimes, including amplified diel variability, persistent hypoxia, and near-anoxia that can precede tissue disintegration and mortality.

**Figure 1.**
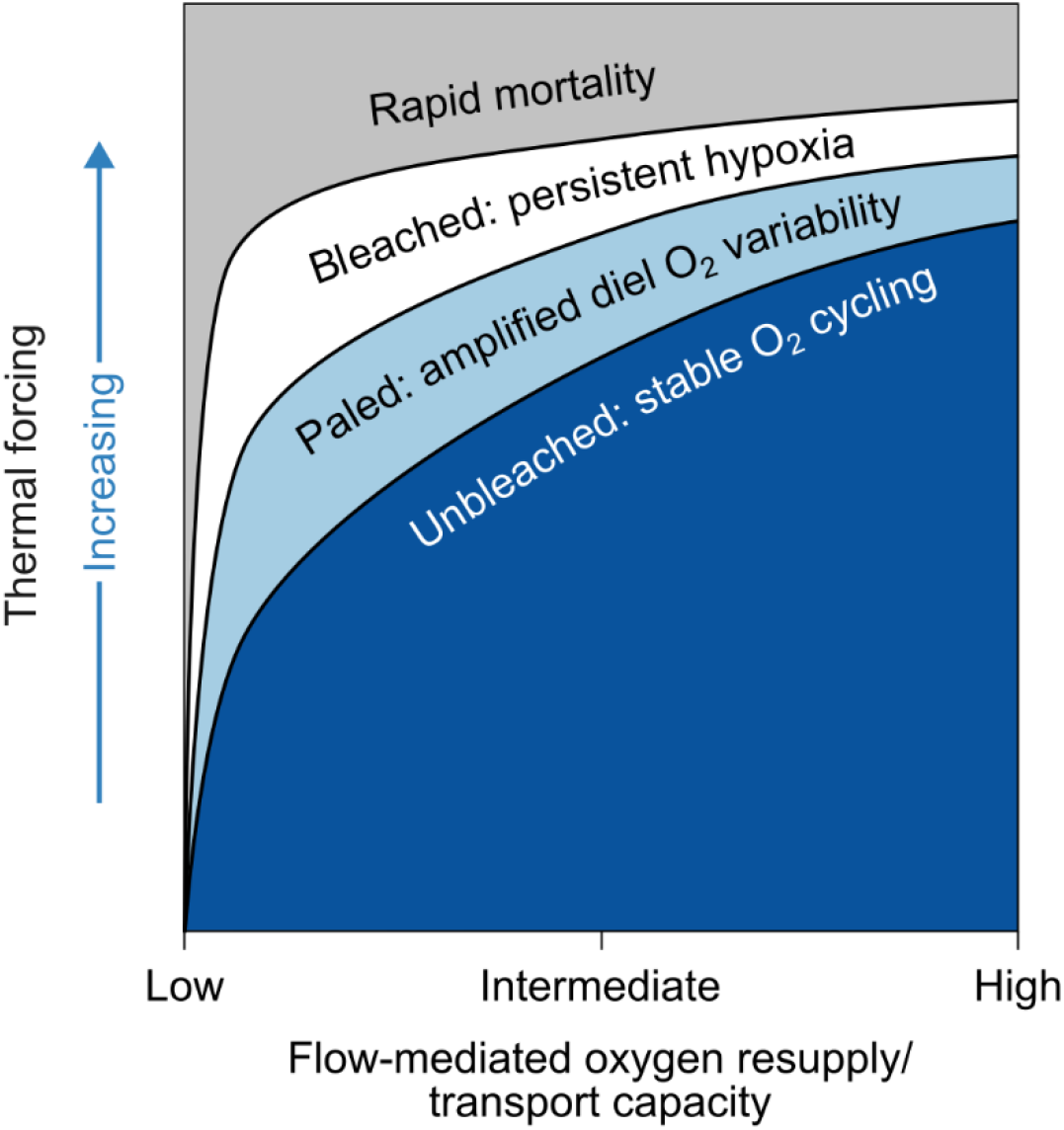
Conceptual transport-metabolism framework for oxygen-driven coral bleaching. Conceptual phase space illustrating coral oxygen microenvironment stability as a function of temperature (y-axis, relative to the Maximum Monthly Mean, MMM) and water flow velocity (x-axis). Colour gradient indicates coral oxygen microenvironment from high (dark blue) to low (grey). Four regimes are delineated by threshold-like boundaries: a stable normoxic regime associated with unbleached colonies (bottom right), an intermediate stress regime associated with paling (centre), a persistent, deoxygenated regime associated with bleaching (upper centre), and an anoxic regime associated with rapid mortality (top).

We tested this framework by integrating high-frequency *in situ* measurements of dissolved oxygen within and around colonies of the branching coral *Pocillopora favosa* during two consecutive Red Sea heatwaves (2023 and 2024) with a three-dimensional model resolving colony-scale flow, oxygen transport, and metabolism. The model, which incorporated co-located measurements of temperature, light, bulk water oxygen, and flow, in a coral reef in the central Red Sea reproduced divergent oxygen regimes and bleaching trajectories consistent with patchy bleaching observed in 2023, and reproduced widespread nighttime near-anoxia associated with the more prolonged and intense 2024 marine heatwave that preceded tissue disintegration and rapid mortality across all coral colonies.

### Contrasting heatwaves reveal divergent and synchronous oxygen instability

During summer-autumn 2023, the central Red Sea experienced an extreme marine heatwave that imposed sustained thermal stress at the study site (Al Fahal reef complex, Fig 2A, B). *In situ* temperatures at 6.5-8 m depth reached daily mean values up to 32.57 °C, with cumulative thermal stress reaching an unprecedented 16.58 degree heating weeks (DHWs, °C-weeks; Fig. 2E, Supplementary Table S1). Within the monitored reef patch (∼2 x 2 m), conspecific *Pocillopora favosa* colonies followed divergent visual bleaching trajectories despite shared environmental conditions. Some colonies remained visually unbleached, whereas others first developed paling within interior colony regions, progressed to whole-colony bleaching, and subsequently recovered (Supplementary Fig. S1; *sensu* ^34^).

**Figure 2.**
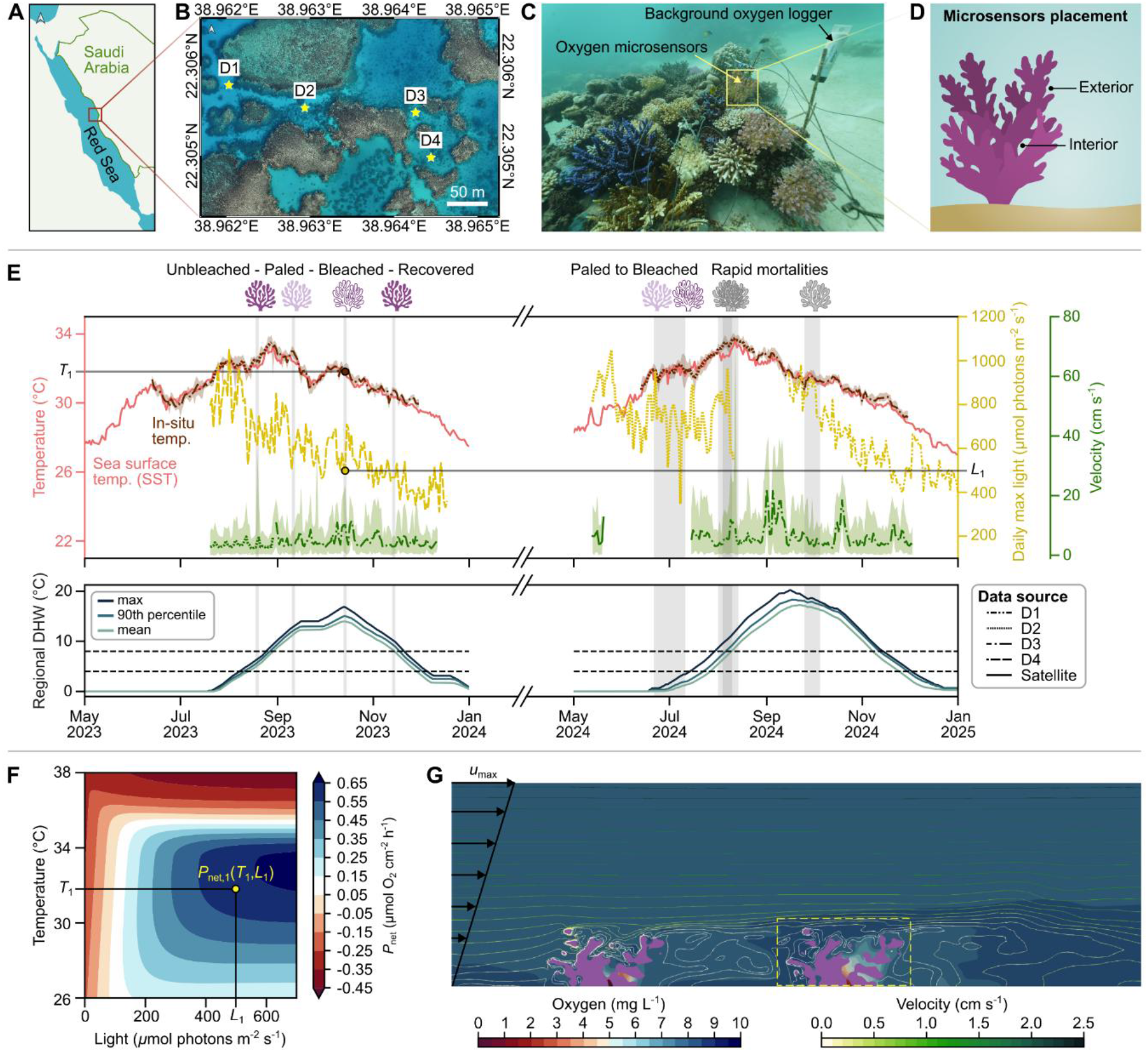
Field site, environmental conditions, and measurement-model framework. **(A-B)** Location of the study site within the Al Fahal reef complex, central Red Sea, Saudi Arabia (22.18°N, 38.96°E), with the monitored reef patch (D2) indicated within the wider reef structure. **(C)** Underwater photograph of the study site showing sensor deployment, with oxygen microsensors positioned within a *Pocillopora favosa* colony and a background oxygen logger deployed in the adjacent water column. **(D)** Schematic of sensor placement showing interior (interbranch) and exterior measurement positions within a branching coral colony. **(E)** Environmental conditions across both heatwave seasons (2023 and 2024), including in situ temperature (black), satellite sea surface temperature (red), daily maximum photosynthetically active radiation (yellow), and flow velocity (green). Coral icons indicate the bleaching state of monitored colonies across deployment periods. Grey shading indicates periods corresponding to Fig. 3 and Supplementary Fig. S5. Lower panel shows Degree Heating Weeks (DHW); dashed lines indicate alert thresholds at 4 and 8 °C-weeks. **(F)** Modelled net photosynthetic oxygen production (P_net_) as a function of light and temperature ^19, 37, 38^; marker indicates representative heatwave conditions (T_1_, L_1_) from panel **(E). (G)** A two-dimensional cutaway of the three-dimensional computational fluid-metabolism model showing the instantaneous flow velocity and dissolved oxygen concentration field for October 24, 2023 at 1600h. The incoming velocity (left) is assumed as a linear profile from 0 to 2.5 cm s^-1^ that flows over two colony geometries, with interior oxygen depletion visible within the colonies.

To characterize oxygen conditions associated with these trajectories, we deployed fine-scale optical oxygen data loggers (AquapHOx-L-O_2_, Pyroscience GmbH) equipped with robust oxygen probes (4mm in diameter). Oxygen was recorded at 1-min intervals within and outside four focal colonies and in the surrounding reef water across four 14-21 day deployments from August to November 2023 (Supplementary Fig. S2).

Despite heatwave conditions, background oxygen remained normoxic throughout all deployments (Fig. 3, Supplementary Fig. S2). Across the representative 48-h intervals, mean dissolved oxygen concentrations ranged from 6.64 to 6.99 mg O_2_ L^−1^, and nighttime minima remained above reported coral hypoxia thresholds of <1-4 mg O_2_ L^−1^ ^30, 35, 36^ (Supplementary Table S2).

**Figure 3.**
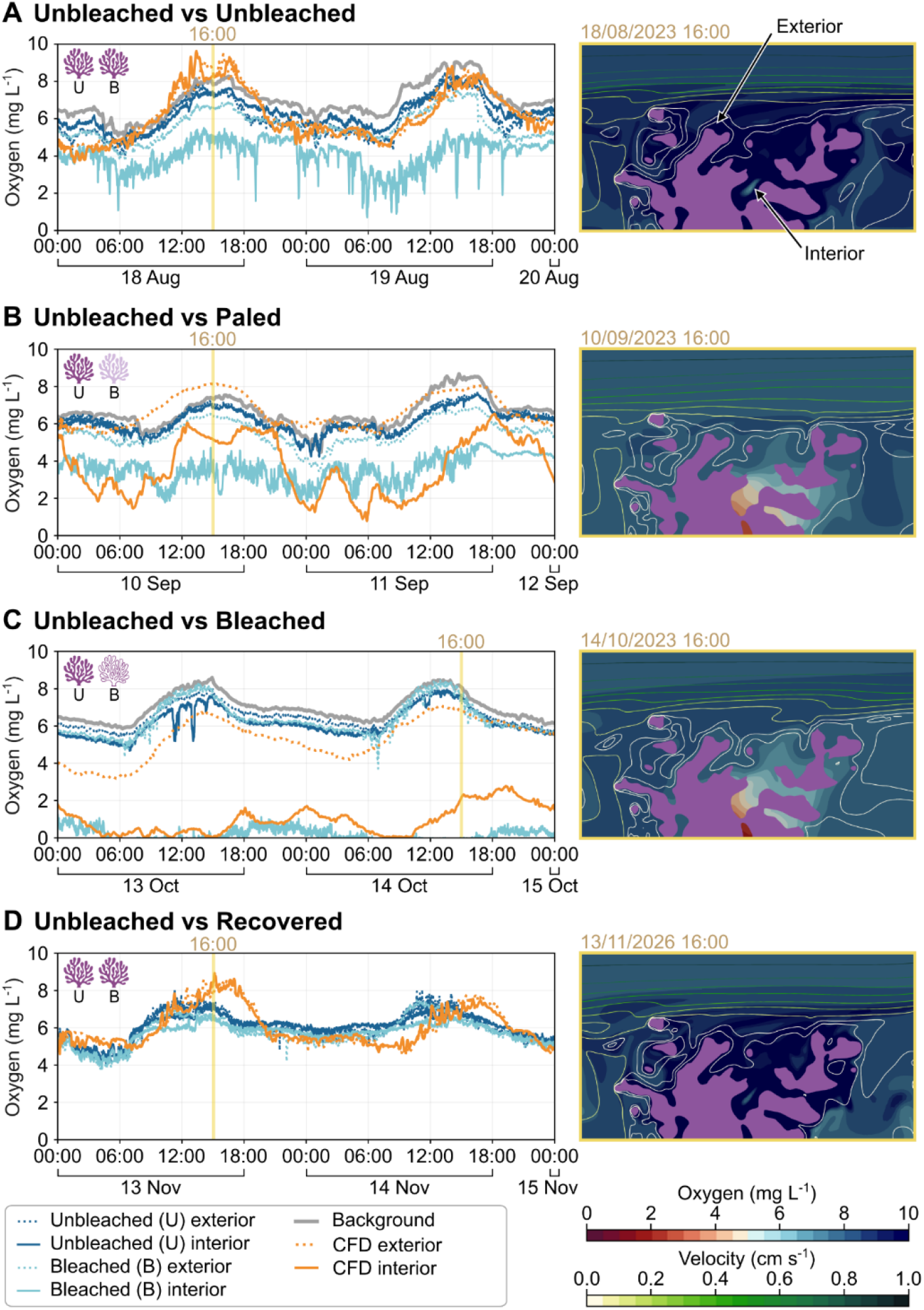
Divergent oxygen microenvironments within coral colonies across the 2023 marine heatwave. Dissolved oxygen time series measured over representative 2-day windows (left panels) and corresponding modelled oxygen microenvironments (right panels) across four successive deployment periods: **(A)** August -unbleached, **(B)** September - paled, **(C)** October - bleached, **(D)** November - recovered. Time series show oxygen concentrations (mg L^-1^) in background reef water (grey), unbleached colony exterior (dark blue dotted) and interior (dark blue solid), bleached colony exterior (light blue dotted) and interior (light blue solid). Orange lines indicate computational fluid-metabolism dynamic model (CFD) output for exterior (dotted) and interior (solid) positions as marked in the right panel of **(A)** for all cases. Right panels show coral cross-sections with simulated oxygen distributions surrounding the coral skeleton (purple), illustrating oxygenated conditions (August, November) and progressive development of hypoxic and near-anoxic microenvironments (September, October), consistent with in situ observations. Yellow vertical lines indicate the time points at which cross-sectional snapshots were extracted. In August and November, interior oxygen concentrations in both colonies track background diel variability. In September, the interior of the bleaching colony exhibits deeper nighttime oxygen minima relative to the unbleached colony. In October, the bleached colony interior collapses to near-anoxic concentrations with minimal daytime recovery, whereas the unbleached colony remains normoxic. CFD simulations reproduce the observed interior-exterior oxygen separation and its progression across bleaching states. Simulations were performed at a constant flow velocity of 2.5 cm s^−1^ for all cases, with net photosynthesis (P_net_) parameterized from in situ light and temperature measurements.

Oxygen conditions within colony interiors, however, diverged strongly between colonies displaying divergent visual bleaching trajectories. Colonies that remained unbleached maintained stable interbranch oxygen concentrations across deployments, with temporal means of 6.07-6.32 mg O_2_ L^−1^ (Supplementary Table S2). By contrast, oxygen records associated with the paling-to-bleaching trajectory showed progressive depletion across deployments, with 48h-mean concentrations declining from 4.04 mg O_2_ L^−1^ in August, to 3.53 mg O_2_ L^−1^ in September, and 0.17 mg O_2_ L^−1^ in October (Fig. 3A-C; Supplementary Table S2). Concomitantly, diel oxygen amplitudes declined from 4.58 mg O_2_ L^−1^ to 1.05 mg O_2_ L^−1^, consistent with diminished daytime photosynthetic oxygen production as visible paling and bleaching progressed. Interior concentrations were 1.6-6.2 mg O_2_ L^−1^ lower than corresponding exterior measurements, and paling was first visible within the same interior colony regions (Supplementary Fig. S1C-E). Full deployment records confirmed increasingly severe nighttime minima and longer low-oxygen exposure from August to October (Supplementary Fig. S2).

During peak thermal stress in October 2023, oxygen within the monitored bleaching-trajectory record remained below 2 mg O_2_ L^−1^ throughout the representative 48-h interval, corresponding to 2,880 min of continuous hypoxia exposure with little daytime recovery (Fig. 3C, Supplementary Table S2). Available records from the unbleached colonies did not cross this threshold during the analysed deployments. Following visual recovery in November, recorded oxygen dynamics returned toward normoxic diel variability comparable to those of unbleached colonies (Fig. 3D; Supplementary Table S2). Together, these observations document a transition from stable diel oxygen cycling to persistent interior hypoxia, followed by a return to normoxic variability after visual recovery, despite consistently normoxic surrounding reef water.

In 2024, a second marine heatwave exposed the central Red Sea to earlier, longer and more intense thermal stress compared to 2023. Daily mean seawater temperatures reached 34.06 °C and °C-weeks began accumulating in mid-June, ultimately reaching 20.22 DHWs (Fig. 2E, Supplementary Table S1). All monitored *P. favosa* colonies in 2024 experienced tissue disintegration and rapid mortality (*sensu* ^25^; Supplementary Fig. S3), eliminating the divergent colony responses observed in 2023. Mean current velocity was 5.58 cm s^−1^ in 2024 compared to 4.78 cm s^−1^ in 2023 (Fig. 2E; Supplementary Table S1). Note that these measurements characterize flow outside the colonies and do not directly resolve transport within interbranch spaces or more sheltered parts of the reef patch. Colonies that died earliest were located in relatively sheltered microhabitats that had previously bleached in 2023 (Supplementary Fig. S4), consistent with local hydrodynamic exposure influencing susceptibility.

Like 2023, surrounding reef seawater remained normoxic throughout all deployments (Fig. 4A, Supplementary Fig. S5, S6). Mean background oxygen ranged from 5.59 to 6.86 mg O_2_ L^−1^, with nighttime minima of 2.62-4.75 mg O_2_ L^−1^ across colony-specific observation periods (Supplementary Table S3). In contrast, all four colony interiors experienced recurrent oxygen depletion with minimum concentrations between 0.09-0.47 mg O_2_ L^−1^ (Supplementary Fig. S5, S6). Mean interior oxygen concentrations ranged from 3.17 to 5.90 mg O_2_ L^−1^, compared to 5.11-6.47 mg O_2_ L^−1^ at exterior positions. Interior oxygen depletion was also more persistent: cumulative exposure below 2 mg O_2_ L^−1^ ranged from 897 of 14,385 valid observation minutes to 3,639 of 28,776 valid observation minutes, equivalent to 6.2-12.7% of the colony-specific observation periods (Supplementary Fig. S6, Supplementary Table S3). By contrast, exterior positions accumulated only 1-113 min below 2 mg O_2_ L^−1^, never exceeding 0.8% of their respective observation periods, while background reef water never crossed this threshold (Supplementary Fig. S6, Supplementary Table S3).

**Figure 4.**
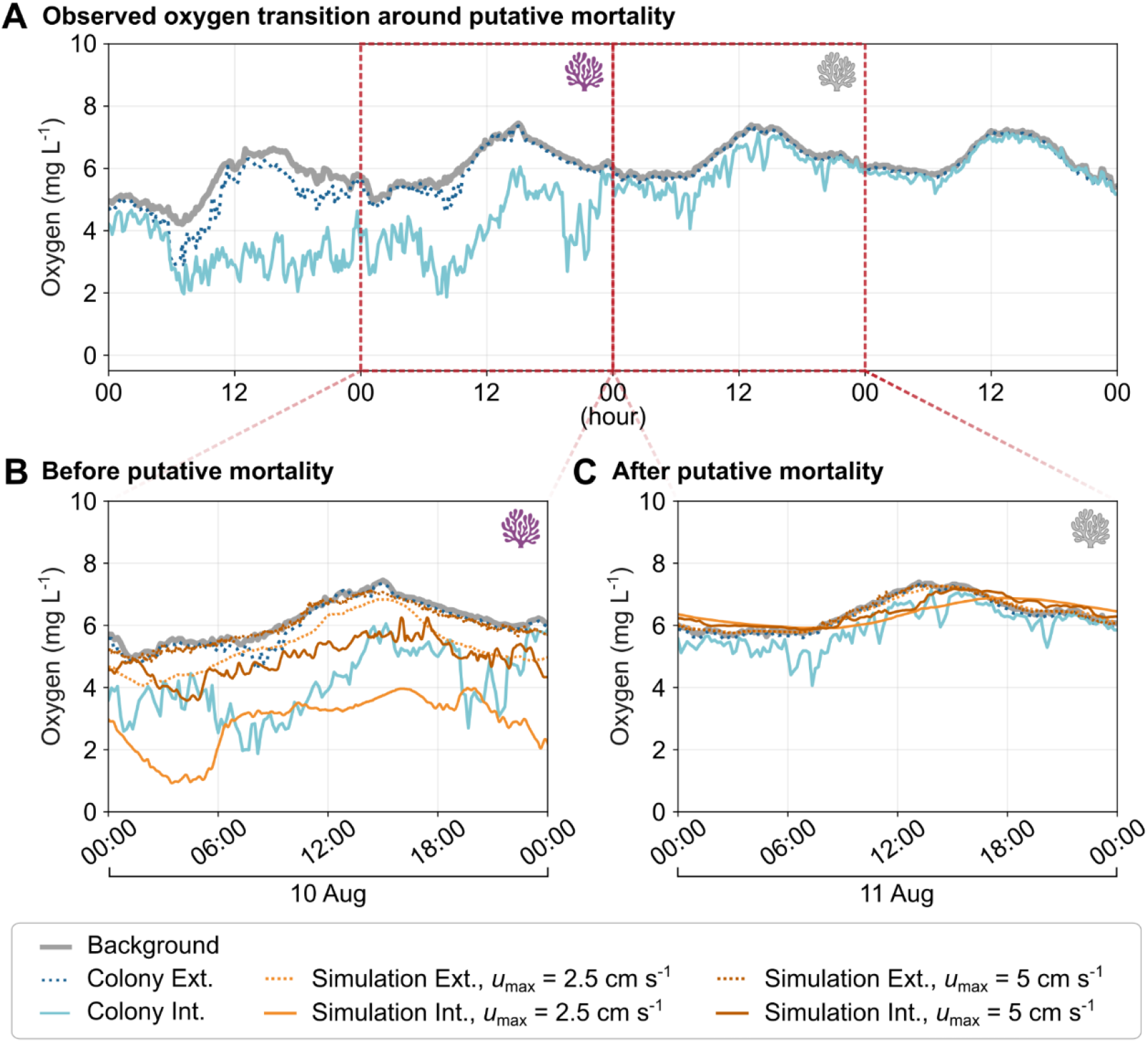
Holobiont metabolism and local flow shape colony oxygen dynamics around putative mortality during the 2024 marine heatwave. **(A)** Field observations from Colony 2 across four diel cycles from 9-12 August 2024, showing dissolved oxygen concentrations in background reef water (grey), at the colony exterior (dotted blue), and within the colony interior (cyan). Outlined windows indicate the 24-h periods enlarged in panels B and C. Coral symbols denote periods before and after putative mortality. **(B)** Field observations before putative mortality compared with simulations retaining holobiont metabolic oxygen exchange at maximum inlet velocities of 2.5 and 5 cm s^−1^. **(C)** Field observations during the corresponding period after putative mortality compared with simulations in which holobiont metabolic oxygen exchange was set to zero at the same inlet velocities. Simulated exterior and interior concentrations are shown as dotted and solid orange lines, respectively; lighter and darker shades denote *u*_max_ = 2.5 and 5 cm s^−1^. In the simulations, active holobiont metabolism generated pronounced interior–exterior separation that increased under lower local flow, whereas setting metabolic oxygen exchange to zero substantially reduced this separation. The simulations were not fitted to the field time series and test the qualitative effects of metabolism and local transport rather than point-by-point agreement.

Unlike the persistent diel damping observed during advanced bleaching in 2023, colonies in 2024 generally retained pronounced diel oscillations while repeatedly experiencing near-anoxia at night. Interior diel amplitudes ranged from 1.83 to 7.99 mg O_2_ L^−1^ across colonies, indicating substantial daytime reoxygenation followed by strong nighttime depletion (Supplementary Table S3). Recurrent nighttime near-anoxia emerged days to weeks before visible tissue loss across all four colonies (Supplementary Fig. S5, S6), although its onset and persistence varied, with some colonies exhibiting sustained oxygen depletion and others intermittent episodes followed by partial daytime recovery (Supplementary Fig. S5). Thus, the stronger thermal stress of 2024 was associated with recurrent interior oxygen limitation across all monitored colonies before tissue loss and mortality, despite normoxic surrounding water and continued daytime oxygen resupply.

### A transport-metabolism model reproduces oxygen regimes across both heatwaves

To determine whether these contrasting oxygen regimes observed across both heatwaves could arise from fundamental principles of transport and metabolism, we developed a three-dimensional fluid-metabolism model that resolves flow, temperature, and oxygen transport within a realistic colony geometry derived (Supplementary Fig. S7) from micro-computed tomography (CT) data. For the 2023 simulations, the model was forced with *in situ* environmental measurements of temperature, light, background oxygen, and flow (Fig. 2E, Supplementary Fig. S8), together with literature-based coral metabolic parameterisations ^19, 37, 38^. This provided an independent test of whether the observed oxygen regimes could emerge from general transport-metabolism interactions, rather than colony-specific physiological differences. Importantly, no parameters were fitted to the observed colony outcomes.

The 2023 simulations reproduced the defining transition from stable diel oxygen variability to persistent depletion within colony interiors (Fig. 3A-C). Reductions in flow to 2.5 cm s^−1^ and net photosynthetic production produced more profound nighttime minima and extended periods of low oxygen availability within interbranch spaces, reproducing intermediate oxygen regimes observed during paling (Fig. 3B). Under conditions representing colony-wide bleaching, simulated interior oxygen remained persistently low with little daytime recovery, despite normoxic boundary conditions (Fig. 3C, Supplementary Table S2). Under conditions representing the stable oxygen regime, including higher model boundary flow (5 cm s^−1^) and intact net photosynthetic oxygen production, simulated interior and exterior oxygen dynamics were consistent with the *in situ* observations (Supplementary Fig. S9).

Model diagnostics demonstrated that oxygen depletion intensified as net metabolic oxygen consumption exceeded transport-mediated resupply for progressively longer periods of each diel cycle, eventually preventing full daytime reoxygenation and allowing oxygen deficits to accumulate within confined interbranch spaces. Time-resolved simulations further illustrate the progression from stable diel cycling to oxygen collapse across bleaching states (Supplementary Videos S1-S4). Under the geometry and environmental forcing tested, reducing flow to approximately 2.5 cm s^−1^ shifted colony interiors from stable oxygen availability to persistent hypoxia. This transition represents a specific outcome of the tested model configuration rather than a general flow threshold.

To determine whether the same mechanism could generate field-like oxygen dynamics under different local transport conditions in 2024, we conducted four simulations under representative high-temperature conditions. The simulations crossed inlet velocities of 2.5 and 5 cm s^−1^ with coral holobiont metabolic oxygen exchange either retained or set to zero. The latter represented an idealized post-mortality state in which photosynthetic oxygen production and consumption by the coral holobiont has ceased (Fig. 4B, C).

At 2.5 cm s^−1^ with holobiont metabolism retained, simulated interior oxygen ranged from 0.92 to 5.37 mg O_2_ L^−1^ and remained, on average, 2.41 mg O_2_ L^−1^ below exterior concentrations. Interior oxygen remained below 2 mg O_2_ L^−1^ for 722 min across the 72-h simulation, whereas exterior oxygen remained above 4.06 mg O_2_ L^−1^. The model thereby generated oxygen dynamics qualitatively consistent with the defining *in situ* pattern of pronounced diel interior-exterior separation, nighttime oxygen depletion, and daytime reoxygenation despite normoxic surrounding water. Increasing inlet velocity to 5 cm s^−1^ reduced the mean interior-exterior difference to 0.95 mg O_2_ L^−1^, maintained interior concentrations above 3.58 mg O_2_ L^−1^, and prevented hypoxia (Fig. 4B).

When holobiont metabolic oxygen exchange was set to zero to represent coral mortality, mean interior and exterior concentrations differed by only 0.03-0.04 mg O_2_ L^−1^ at either inlet velocity, and no hypoxia developed. Interior oxygen continued to follow variation in exterior oxygen with a temporal delay, but this delay decreased at the higher inlet velocity, consistent with faster convective transport (Fig. 4C). Thus, colony geometry and restricted flow alone were insufficient to generate sustained interior oxygen depletion. Field-like oxygen instability emerged from the interaction between metabolic oxygen exchange and restricted transport.

Together, the same transport-metabolism model generated the contrasting oxygen signatures observed across both heatwaves.

### Oxygen instability links heatwave intensity to coral fate

Heatwave exposure is often summarized at reef-to-regional scales, yet our measurements show that neighbouring coral colonies concurrently experienced distinct oxygen microenvironments during the same event. Across two consecutive marine heatwaves, intact branching coral colonies developed low-oxygen microenvironments before visible paling, bleaching, tissue disintegration, or mortality, despite consistently oxygenated surrounding seawater. By combining high-frequency *in situ* observations with a three-dimensional transport-metabolism model, we provide independent evidence supporting an imbalance between temperature-dependent metabolic oxygen demand and flow-mediated oxygen resupply as a mechanism leading to this instability.

The two heatwaves occurred during the fourth global bleaching event, when marine heatwaves generated extreme accumulated heat stress across most reef regions ^1, 6^. The resulting impacts were severe, with initial estimates suggesting that 43% of affected reefs experienced substantial mortality during 2023-2024, compared with 15% during the 2014-2017 global bleaching event ^1, 39^. Thermal stress reached 16.58 and 20.22 DHW in 2023 and 2024, respectively, accumulating earlier and reaching higher temperatures in 2024 at the study area ^7^. In 2023, neighbouring colonies exposed to shared bulk-water conditions developed oxygen regimes aligned with their subsequent fates: stable diel cycling accompanied the unbleached trajectory, whereas progressive oxygen depletion preceded paling, bleaching before recovery. In 2024, this divergence gave way to recurrent near-anoxia before tissue disintegration and rapid mortality across all monitored colonies, despite far-field flow velocities and background oxygen concentrations comparable to or greater than those measured in 2023. Similar rapid tissue disintegration was reported during extreme heat stress in the Caribbean, where dominant reef-building corals exhibited tissue ‘sloughing’ rather than conventional bleaching ^40^. These observations suggest that sufficiently intense heatwaves compress or bypass the prolonged progression from paling to bleaching and instead lead directly to rapid mortality ^25^. Within the framework illustrated in Fig. 1, 2023 therefore reflects a transport-sensitive regime in which local hydrodynamic differences may have helped separate colony fates. By contrast, the 2024 field observations reflect a thermally overwhelmed regime in which elevated metabolic demand with stronger heat stress exceeded the buffering capacity of local transport and drove widespread oxygen instability and mortality. The high-flow simulations show that this response is conditional on local transport, which is not captured by external mean velocity alone.

Our findings extend the compound heat-oxygen framework from ecosystem to organism scales. Marine heatwaves increasingly coincide with deoxygenation because warming elevates biological oxygen demand, reduces oxygen solubility, and can restrict ventilation through altered mixing or flow ^41–45^. Our results show that the same physical-biological imbalance can also arise within individual organisms, whereby thermally altered metabolism generates localized oxygen deficits in flow-restricted interbranch spaces while the surrounding reef water remains normoxic. Oxygen exposure within a coral colony therefore cannot be inferred from bulk-water oxygen concentration measurements alone, but depends on the interaction among thermal regime, metabolic state, colony architecture, and local hydrodynamic exchange.

Flow regulates the exchange of oxygen and other solutes between colony interiors and surrounding water. Branching morphologies can restrict advective and diffusive transport, allowing oxygen consumption to outpace replenishment when metabolic demand is high ^26–28, 46^. This framework provides a basis for earlier observations that elevated water flow can reduce coral mortality during thermal stress and facilitate bleaching recovery ^31, 32^. Although the simulations support this interpretation, the zero-metabolism endpoint is intentionally simplified. Recently dead coral skeletons can undergo rapid, microbially mediated decay following heatwave mortality ^47^. Such post-mortality metabolism was not represented in the model and may contribute to differences between simulated and observed oxygen dynamics following tissue loss. Flow should therefore be interpreted as a context-dependent buffer rather than a universal velocity threshold. The temporal structure of oxygen exposure differed between coral fates: sustained hypoxia with little daytime recovery characterized the bleaching trajectory in 2023, whereas colonies in 2024 repeatedly approached near-anoxia at night but generally reoxygenated during the day before tissue disintegration. Repeated nighttime oxygen deficits may accumulate stress across successive days ^48, 49^, particularly under sustained thermal stress ^50^. However, overlapping minimum oxygen concentrations between bleaching in 2023 and mortality in 2024 indicate that the present observations do not define fixed oxygen thresholds. Mortality may instead reflect lower minima, greater cumulative oxygen deficits, faster thermal stress accumulation, or interactions among these factors.

Marine heatwaves remain the ultimate driver of mass bleaching and mortality, but our findings identify organism-scale oxygen imbalance as a mechanism connecting shared thermal exposure to local response variability and, eventually, collapse. Incorporating oxygen exposure and local hydrodynamic exchange into reef monitoring and modelling may therefore improve predictions of when shared thermal extremes drive divergent coral fates and guide a more climate-resilient deployment of coral restoration efforts.

## Acknowledgements

We are thankful for members of the Marine Microbiomes Lab, especially Dr. Matteo Monti, Dr. Erika P. Santoro and Mr. Marco Casartelli, for their help during fieldwork and photograph editing, and to the Coastal and Marine Resources Core lab for their continuous support, including diving operations, oceanographic equipment, marine permits, and operations. We thank Dr. Anieka Parry and co-authors for allowing early access to data prior to their publication which informed parameterisation of the model generated here. We gratefully acknowledge Prof. Thomas Finkbeiner and Dr. Nikolay Gorshkov at the GEMs Platform for performing the CT scan and generating the digital model of the specimen.

## Author contributions

Conceptualization: YEK, SGK, TTT, HHK, NGB, FCG, ST, and MP; Data curation: YE, FCG, and NGB; Experimental investigation: YEK, NGB, SGK, TTT, and ST; Model parameterization & input variable determination: YEK, SGK, TTT, MP, HHK, MSD, ST, DM, AKK, and SD; Modelling and software development: HHK, TM, MP, DM, AKK, and SD; Model analysis and validation: HHK, MP, MO, DM, AKK, SD, and TTT; Visualization: YEK, HHK, TTT, FW, NGB, and SGK; Funding acquisition: TTT, MP, RSP, MJ, MF, CMD, and SC; Supervision: TTT, MP, YEK, MJ, MF, SC, and RSP; Writing – original draft: YEK; Writing – review & editing: all

## Data availability

Correspondence and requests for supplementary materials and related information should be addressed to Dr. Yusuf C. El-Khaled and Prof. Tadd T. Truscott.

